# Trafficking regulator of GLUT4-1 (TRARG1) is a GSK3 substrate

**DOI:** 10.1101/2020.12.07.415257

**Authors:** Xiaowen Duan, Dougall M. Norris, Sean J. Humphrey, Pengyi Yang, Kristen C. Cooke, Will P. Bultitude, Benjamin L. Parker, Olivia J. Conway, James G. Burchfield, James R. Krycer, Frances M. Brodsky, David E. James, Daniel J. Fazakerley

**Affiliations:** Charles Perkins Centre, School of Life and Environmental Sciences, The University of Sydney, Sydney, NSW 2006, Australia; Metabolic Research Laboratories, Wellcome-Medical Research Council Institute of Metabolic Science, University of Cambridge, Cambridge, CB2 0QQ, UK; Charles Perkins Centre, School of Mathematics and Statistics, The University of Sydney, Sydney, NSW 2006, Australia; Research Department of Structural and Molecular Biology, Division of Biosciences, University College London, Gower Street, London WC1E 6BT, UK; School of Medicine, The University of Sydney, Sydney, NSW 2006, Australia; Computational Systems Biology Group, Children’s Medical Research Institute, The University of Sydney, Westmead, NSW 2006, Australia

**Keywords:** glycogen synthase kinase 3 (GSK3), glucose transporter type 4 (GLUT4), trafficking regulator of GLUT4-1 (TRARG1), protein phosphorylation, phosphoproteomics, insulin signaling

## Abstract

Trafficking regulator of GLUT4-1, TRARG1, positively regulates insulin-stimulated GLUT4 trafficking and insulin sensitivity. However, the mechanism(s) by which this occurs remain(s) unclear. Using biochemical and mass spectrometry analyses we found that TRARG1 is dephosphorylated in response to insulin in a PI3K/Akt-dependent manner and is a novel substrate for GSK3. Priming phosphorylation of murine TRARG1 at serine 84 allows for GSK3-directed phosphorylation at serines 72, 76 and 80. A similar pattern of phosphorylation was observed in human TRARG1, suggesting that our findings are translatable to human TRARG1. Pharmacological inhibition of GSK3 increased cell surface GLUT4 in cells stimulated with a submaximal insulin dose, and this was impaired following *Trarg1* knockdown, suggesting that TRARG1 acts as a GSK3-mediated regulator in GLUT4 trafficking. These data place TRARG1 within the insulin signaling network and provide insights into how GSK3 regulates GLUT4 trafficking in adipocytes.

## Introduction

Insulin maintains whole-body glucose homeostasis by regulating glucose transport into muscle and adipose tissue and by inhibiting glucose output from liver. Defects in these processes cause insulin resistance, which is a precursor to metabolic disorders including Type 2 diabetes. Insulin signaling augments glucose transport into adipose tissue and muscle by promoting the translocation of the glucose transporter GLUT4 from a specialized intracellular storage compartment (GLUT4 storage vesicles; GSVs) to the plasma membrane (PM) [1]. Previous proteome studies of intracellular membranes highly enriched in GLUT4 have revealed major GSV resident proteins including TBC1D4 (AS160), Sortilin, IRAP, LRP1, VAMP2 [2–4] and Trafficking regulator of GLUT4-1 (TRARG1). TRARG1 positively regulated insulin-stimulated glucose transport and GLUT4 trafficking, and was required for the insulin-sensitizing action of the PPARγ agonist rosiglitazone [5, 6]. However, the mechanisms by which insulin signals to TRARG1 or how TRARG1 promotes insulin sensitivity remains unknown.

The most characterized signaling pathway relevant to GLUT4 trafficking is the PI3K/AKT axis, which is activated by a series of upstream signaling events triggered by the binding of insulin to its receptor at the cell surface. AKT is necessary and sufficient for insulin-stimulated GLUT4 trafficking [7], and is thought to act predominantly via phosphorylation of the Rab GTPase-activating protein (GAP), TBC1D4. Phosphorylation of TBC1D4 inhibits its GAP activity, and permits its target Rab GTPases to mediate the translocation of GSVs to the PM [8]. However, loss of TBC1D4 or its cognate Rab proteins did not completely mimic or inhibit insulin-stimulated GLUT4 translocation [9–11], suggesting that there may be additional sites of insulin action within the GLUT4 trafficking pathway [12].

In addition to phosphorylating substrates such as TBC1D4, AKT also modulates cellular metabolism through increased or diminished activity of other kinases. For example, AKT directly phosphorylates glycogen synthase kinase 3 (GSK3) at the regulatory Ser21 or 9 residue (α and β isoform, respectively), which inhibits its kinase activity [13] and leads to dephosphorylation and activation of glycogen synthase (GS) and thereby glycogen synthesis. GSK3 is a unique kinase in that 1) it is constitutively active and inhibited in response to extracellular stimulation and 2) its substrates usually need priming by another kinase [14]. Given the role of GSK3 in glycogen synthesis, it has generally been thought that GSK3 mainly participates in glucose metabolism via regulation of glycogen synthesis [15–19]. However, although there are conflicting reports [20–22], evidence from studies using small molecules to acutely inactivate GSK3 shows that GSK3 may also regulate glucose transport and GLUT4 translocation [21, 22].

In the present study, we identified a range of post-translational modifications (PTMs) on TRARG1 and integrated TRARG1 into the insulin signaling pathway by showing that insulin promotes TRARG1 dephosphorylation via PI3K/AKT. Specifically, we report that TRARG1 is a novel direct substrate of the protein kinase GSK3, which phosphorylates murine TRARG1 at three sites within a highly phosphorylated patch between residues 70 and 90 in the cytosolic portion of TRARG1. Further, submaximal insulin-stimulated GLUT4 trafficking was potentiated by GSK3 inhibition, but this potentiation was impaired by simultaneous *Trarg1* knockdown. Overall, our findings have revealed new information on how TRARG1 is regulated by insulin signalling and suggest that TRARG1 may provide a link between GSK3 and GLUT4 trafficking.

## Results

### TRARG1 phosphorylation alters its migration in SDS-PAGE

We previously reported that a proportion of TRARG1 exhibited reduced electrophoretic mobility following separation by SDS-PAGE. These apparent higher molecular weight species of TRARG1 were enriched in the PM but not low or high density microsomal (LDM, HDM) fractions (Fig. 1A) [5]. Given the role of TRARG1 in GLUT4 trafficking, we sought to identify the cause of altered electrophoretic mobility of TRARG1 with the aim of providing insight into how TRARG1 regulates GLUT4. We hypothesized that these apparent higher molecular weight TRARG1 bands are due to PTM of TRARG1 as: 1) they were all reduced in intensity upon knock-down of TRARG1 using different sets of siRNAs [5], and 2) they are not splice variants as the cDNA of TRARG1, which does not contain any introns, also generates multiple bands when transfected into HEK-293E cells (Fig. 1B).

**Figure 1.**
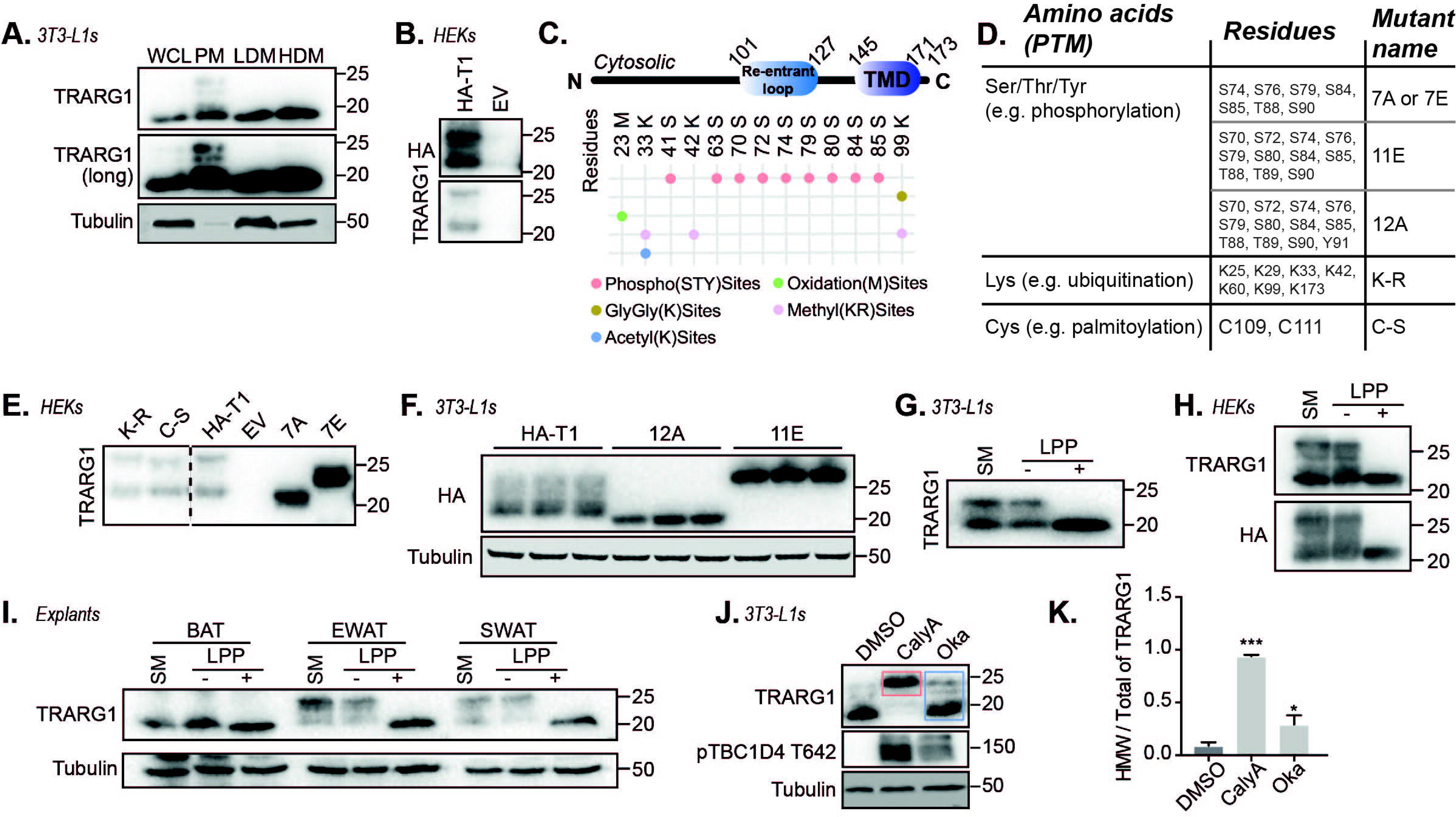
TRARG1 phosphorylation causes apparent higher molecular weight bands by immunoblotting. (A) Subcellular fractionation of 3T3-L1 adipocytes. A longer exposure time for the TRARG1 blot is presented to better visualize higher molecular weight bands (TRARG1 (long)). Apparent higher molecular weight TRARG1 bands are enriched in PM fractions (WCL; whole cell lysate, PM; plasma membrane, LDM; low density microsomes, HDM; high densty microsomes). (B) HA-tagged murine TRARG1 (HA-TRARG1) expressed in HEK-293E cells shows multiple bands by immunoblotting (HA-T1; N-terminally tagged HA-TRARG1, EV; empty vector control). (C) Schematic of the domains in TRARG1 and post-translational modifications detected by mass spectrometry on HA-TRARG1 (murine) overexpressed in HEK-293E cells. (D) Table of TRARG1 Ser/Thr/Tyr, Lys and Cys mutants used to study TRARG1 post-translocation modifications (PTMs) in Fig. 1. Murine TRARG1 residue positions are indicated. (E) HA-TRARG1 phosphomutants with Ser/Thr mutated to Ala or Glu (7A/7E) expressed in HEK-293E cells exhibited a molecular weight similar to the apparent lower or higher molecular weight of HA-TRARG1, respectively. Lys-Arg (K-R) and Cys-Ser mutation had no effect on TRARG1 band patterning. (F) Immunoblotting analysis of TRARG1 phosphomutants (12E, 11A) expressed in 3T3-L1 adipocytes. (G) *In vitro* Lambda protein phosphatase (LPP) treatment of 3T3-L1 adipocytes lysates removed apparent higher molecular weight TRARG1 bands. (H) Apparent higher molecular weight TRARG1 bands were removed by LPP treatment in TRARG1 expressed in HEK-293E cells. (I) Apparent higher molecular weight TRARG1 bands were present in white adipose tissue (epididymal white adipose tissue; EWAT, subcutaneous white adipose tissue; SWAT) lysates, but not brown adipose tissue (BAT) lysates as analyzed by immunoblotting. The apparent higher molecular weight TRARG1 bands were removed by LPP treatment. (J) Apparent higher molecular weight TRARG1 bands were increased in intensity in 3T3-L1 adipocytes treated with phosphatase inhibitors, calyculin A (CalyA) or okadaic acid (Oka). (K) Quantification of (J). The ratio of apparent higher molecular weight (HWM) TRARG1 (as indicated by the bands in the *pink* box in (J)) signal to total TRARG1 (as indicated by the bands in the *blue* box in (J)) signal was quantified as a metric of TRARG1 phosphorylation (n=3, mean±SEM, \**p* <0.05; \*\*\**p* <0.001, comparisons with cells under DMSO condition). For panels A, B, D, E, F, G, H and I, the migration positions of molecular mass markers (kilodaltons) are shown to the right.

We first used mass spectrometry (MS) analysis to identify PTMs on immunoprecipitated HA-tagged TRARG1 (HA-TRARG1) expressed in HEK-293E cells. This revealed multiple PTM types on murine TRARG1 including phosphorylation, methylation, oxidation, ubiquitination (or other ubiquitin-like modifications as indicated by peptides containing di-gly modifications) and acetylation (Fig. 1C, Supplemental table S1). This analysis revealed TRARG1 to be extensively post-translationally modified, with a particularly high number of phosphorylated sites within a short cytosolic region between residues 70 and 90. To test if these specific PTMs affected TRARG1 mobility in SDS-PAGE, we generated a series of HA-TRARG1 mutants (Fig. 1D), including Ser/Thr to Ala or Glu mutants where Ala prevents and Glu mimics phosphorylation [23], a Lys-null mutant (K→R) to prevent ubiquitination or acetylation of Lys and a Cys-null mutant (C→S) to prevent palmitoylation of Cys. Of these mutants, only the Ser/Thr to Ala mutant expressed in HEK-293E cells or 3T3-L1 adipocytes exhibited a similar electrophoretic mobility as the lower band of HA-TRARG1, while Ser/Thr to Glu mutant migrated to a similar position as the higher bands of HA-TRARG1 (Fig. 1E–F). These data suggest that the apparent higher molecular weight TRARG1 bands are dependent on PTMs on Ser/Thr, but not Lys or Cys.

Consistent with these data, treatment of lysates from 3T3-L1 adipocytes (Fig. 1G), HEK-293E cells transfected with HA-TRARG1 (Fig. 1H) or adipose tissue (Fig. 1I) with Lambda protein phosphatase (LPP) *in vitro* completely removed apparent higher molecular weight TRARG1 bands. We note that the apparent higher molecular weight TRARG1 bands were only present in epididymal and subcutaneous white adipose depots, but not in brown adipose tissue, as previously observed [5] (Fig. 1I). Further, acute treatment of 3T3-L1 adipocytes with the phosphatase inhibitors Calyculin A or Okadaic Acid decreased TRARG1 migration in SDS-PAGE (Fig. 1J–K). Therefore, TRARG1 is extensively post-translationally modified, with phosphorylation reducing TRARG1 mobility by SDS-PAGE. Additionally, the ratio of apparent higher molecular weight TRARG1 signal to total TRARG1 signal by immunoblotting can serve as a proxy metric for TRARG1 phosphorylation (Fig. 1K).

### TRARG1 is dephosphorylated with insulin in a PI3K/AKT-dependent manner

Given the role of TRARG1 in adipocyte insulin responses [5, 6], we next tested whether TRARG1 phosphorylation was regulated by insulin signaling. Using the ratio of apparent higher molecular weight TRARG1 versus total TRARG1 as a proxy for TRARG1 phosphorylation, we detected a decrease in TRARG1 phosphorylation in 3T3-L1 adipocytes stimulated with insulin (Fig.2 A–B). This decrease was blocked by PI3K (wortmannin) or AKT inhibition (MK-2206), but not by mTORC1 (rapamycin) or ERK1/2 inhibition (GDC-0994) (Fig. 2A–B). As evidence for the efficacy of kinase inhibition, phosphorylation of AKT S473 and TBC1D4 T642 was impaired by wortmannin and MK-2206; 4EBP1 S65 by wortmannin, MK-2206 and rapamycin; and p90RSK T359/S363 by GDC-0994 (Fig. 2A). These data indicate that TRARG1 is dephosphorylated in response to insulin in a PI3K/AKT-dependent manner. To complement this, we mined data from a SILAC-based, global phosphoproteomic analysis of insulin-stimulated 3T3-L1 adipocytes previously published by our laboratory [24]. These data revealed five phosphosites on TRARG1 (Ser45, Ser76, Ser79, Ser80 and Ser90) downregulated with insulin treatment compared to basal conditions (log2FC ≤ −0.58) (Fig. 2C). The most insulin-sensitive sites were Ser76, Ser79 and Ser80 (Fig. 2C). AKT inhibition using MK-2206 before insulin stimulation increased the phosphorylation at Ser76 and Ser80 by ≥ 1.5-fold compared to insulin stimulation alone (Fig. 2C). Together these data indicate that phosphosites on TRARG1 are dephosphorylated in response to insulin in a PI3K/AKT-dependent manner and that target sites S76 and S80 appear to be affected.

**Figure 2.**
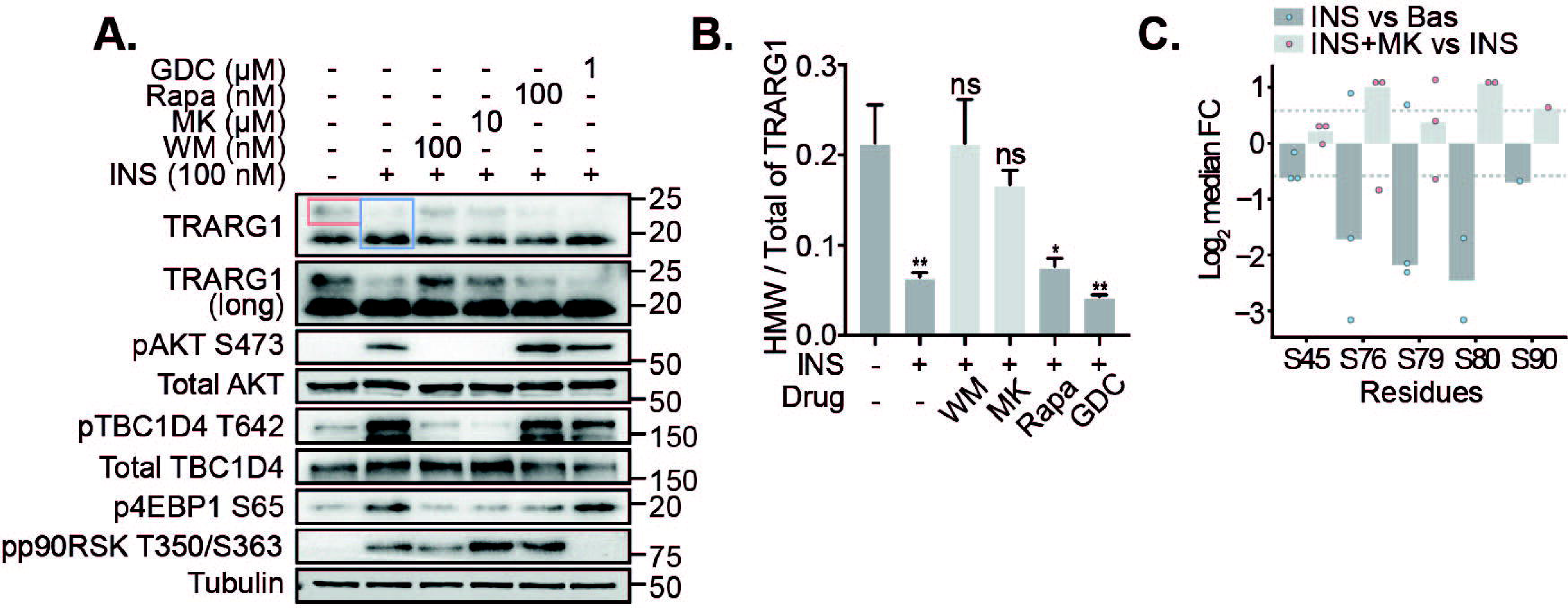
TRARG1 is dephosphorylated with insulin in a PI3K/AKT-dependent manner. (A) 3T3-L1 adipocytes were serum-starved prior to insulin (INS) stimulation (100 nM, 20 min). Where indicated, a PI3K inhibitor (wortmannin (WM), 100 nM), Akt inhibitor (MK-2206 (MK), 10 μM), mTOR inhibitor (rapamycin (rapa), 100 nM), or MAPK inhibitor (GDC-0994 (GDC), 1 μM) was administered 20 min prior to insulin treatment. Samples were analyzed by immunoblotting. A longer exposure time for the TRARG1 blot is presented to better visualize higher molecular weight bands (TRARG1 (long)). (B) Quantification of (A). The ratio of apparent higher molecular weight (HMW) TRARG1 (as indicated by the bands in the *pink* box in (A)) signal to total TRARG1 (as indicated by the bands in the *blue* box in (A)) signal was quantified as a metric of TRARG1 phosphorylation (n=3, mean±SEM, \**p* <0.05; \*\**p* <0.01; ns, non-significant, comparisons with cells without insulin or drug treatment). (C) Bar plot indicating the log_2_-transformed median fold change (FC) in phosphorylation of insulin versus basal or insulin+MK versus insulin alone at Class I TRARG1 phosphosites reported by Humphrey et al. [24]. SILAC-labelled adipocytes were serum-starved prior to insulin stimulation (100 nM, 20 min). The Akt inhibitor, MK-2206 (10 μM), was administered 30 min prior to insulin treatment where indicated. Only sites downregulated (log_2_ FC −0.58) following insulin stimulation are shown. Dashed lines indicate where log_2_ FC =0.58 or −0.58. For panel A, the migration positions of molecular mass markers (kilodaltons) are shown to the right.

### TRARG1 is a novel substrate of GSK3

To elucidate the chain of events by which insulin decreases TRARG1 phosphorylation, we next aimed to identify the kinase responsible for TRARG1 phosphorylation. Here, we considered PKA or GSK3; both kinases that are deactivated by insulin. To test for the involvement of PKA in TRARG1 phosphorylation, we treated 3T3-L1 adipocytes with insulin and/or isoproterenol, a β-adrenergic receptor agonist that potently activates PKA. Insulin-stimulation decreased phosphorylation of both TRARG1 and HSL, a PKA substrate (Fig. 3A). Isoproterenol rescued HSL phosphorylation, but not TRARG1 phosphorylation, suggesting that TRARG1 is not a PKA substrate. In contrast, the GSK3 inhibitors, CHIR99021 and LY2090314, both lowered phosphorylation of glycogen synthase at S641, a GSK3 substrate, and decreased TRARG1 phosphorylation (Fig. 3B). The same effect was observed for endogenous and transfected TRARG1 in six other cell types or tissues. These include endogenous TRARG1 in epididymal (EWAT) and subcutaneous (SWAT) fat explants, and cultured human SGBS adipocytes, treated with insulin or GSK3 inhibitors (Fig. 3C–G), and ectopically expressed TRARG1 in HEK-293E cells, HeLa cells and L6 myotubes treated with GSK3 inhibitors (Fig. 3H–J).

**Figure 3.**
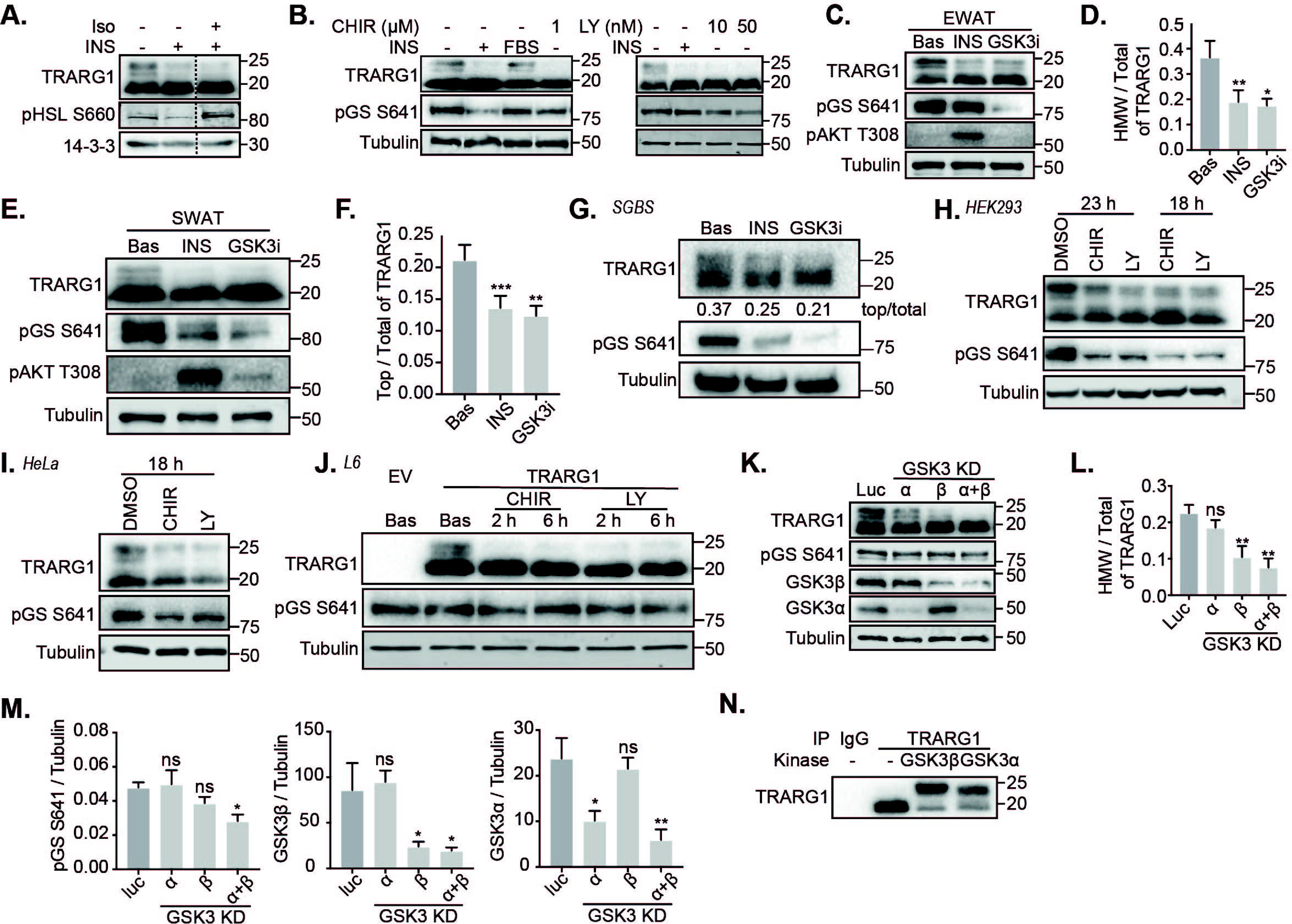
TRARG1 is a GSK3 substrate. (A) 3T3-L1 adipocytes were serum-starved, followed by treatment with insulin (INS; 100 nM) or insulin (100 nM) and isoproterenol (Iso; 1 nM) for 20 min. Samples were analyzed by immunoblotting (Hormone sensitive lipase; HSL). (B) 3T3-L1 adipocytes were serum-starved (except for the *FBS* lane), followed by treatment with insulin (100 nM), CHIR99021 (CHIR) (left panel) or LY2090314 (LY) (right panel) at indicated doses for 20 min. Samples were analyzed by immunoblotting (Glycogen synthase; GS). (C) Epididymal white adipose tissue (EWAT) was excised from mice and minced. Explants were serum-starved in DMEM/2% BSA/20 mM HEPES, pH 7.4 for 2 h followed by treatment with insulin (10 nM) or LY2090314 (GSK3i) (500 nM) for 30 min at 37 °C. Treatment was terminated and tissues were solubilized in RIPA buffer and subjected to analysis by immunoblotting. (D) Quantification of (C). (E) Subcuatneous white adipose tissue (SWAT) was excised from mice and minced. Explants were serum-starved in DMEM/2% BSA/20 mM HEPES, pH 7.4 for 2 h followed by treatment with insulin (10 nM) or LY2090314 (GSK3i) (500 nM) for 30 min at 37 °C. Treatment was terminated and tissues were solubilized in RIPA buffer and subjected to analysis by immunoblotting. Apparent higher molecular weight TRARG1 bands in subcutaneous white adipose tissue (SWAT) explants were reduced in intensity by insulin or GSK3 inhibitor treatment. (F) Quantification of (E). The ratio of apparent higher molecular weight (HMW) TRARG1 signal to total TRARG1 signal was quantified (n=4, mean±SEM, \**p* <0.05; \*\**p* <0.01; \*\*\**p* <0.001, comparisons with basal condition). (G) Apparent higher molecular weight TRARG1 bands in human SGBS adipocytes were reduced in intensity by insulin or GSK3 inhibitor treatment. (H-J) Treatment of HEK-293E cells (H), HeLa cells (I) or L6 myotubes (Empty vector control; EV) (J) with GSK3 inhibitors reduced the intensity of apparent higher molecular weight TRARG1 bands. (K) 3T3-L1 adipocytes on day 6 post-differentiation were transfected with esiRNA targeting GSK3α and/or β. Cells transfected with esiRNA targeting luciferase (Luc) were used as control. 72 h post-transfection, cells were serum-starved for 2 h and harvested for analysis by immunoblotting. (L-M) Quantification of (H). The level of TRARG1 phosphorylation, glycogen synthase (GS) phosphorylation, GSK3α and GSK3β in were quantified (n=3, mean±SEM, \**p* <0.05; \*\**p* <0.01, comparisons with cells transfected with esiRNA targeting luciferase). (N) 3T3-L1 adipocytes were serum-starved in basal DMEM media containing 100 nM LY2090314 for 2 h. Cells were lysed and homogenized and cell lysates were immunoprecipitated with anti-TRARG1 antibody or IgG as a control. Immunoprecipitated TRARG1 was treated with GSK3α, β or reaction buffer alone. All samples were analyzed by immunoblotting. For panels A, B, C, E, G, H, I, J, K and N, the migration positions of molecular mass markers (kilodaltons) are shown to the right.

To complement this pharmacological approach, and to test for a GSK3 isoform-specific effect on TRARG1 phosphorylation, we knocked down *Gsk3*α and/or *Gsk3*β, in 3T3-L1 adipocytes and assessed the phosphorylation status of TRARG1 (Fig. 3K–L). Phosphorylation of glycogen synthase was only significantly decreased with the knock down of both *Gsk3* isoforms (Fig. 3K and M). Phosphorylation of TRARG1 was significantly decreased under conditions where *Gsk3*β was depleted (Fig. 3K–M), whereas knockdown of *Gsk3*α alone did not affect TRARG1 phosphorylation (Fig. 3K–M). These data suggest that TRARG1 is predominantly phosphorylated by GSK3β in 3T3-L1 adipocytes.

We next utilized an *in vitro* assay to test if GSK3 phosphorylates TRARG1 directly. Endogenous TRARG1 was immunoprecipitated from serum-starved 3T3-L1 adipocytes treated with LY2090314 to ensure that the majority of TRARG1 was dephosphorylated while the priming site remained phosphorylated and incubated with recombinant GSK3α or GSK3β. Incubation with either GSK3 isoform decreased TRARG1 electrophoretic mobility (Fig. 3N), suggesting that TRARG1 was phosphorylated by both GSK3α and GSK3β *in vitro*. Together, these data suggest that TRARG1 phosphorylation is regulated by insulin through the PI3K-AKT-GSK3 axis.

### Identification of GSK3 target sites on TRARG1

In general, GSK3 has a substrate consensus motif where it phosphorylates a serine or threonine reside if there is a pre-phosphorylated (primed by a different kinase) serine/threonine four residues C-terminal to the target site (Fig. 4A) [14]. Substrates of GSK3 often contain three or four adjacent S/T-X-X-X-pS/T motifs, allowing GSK3 to phosphorylate every fourth residue in a string of sequential sites as it creates its own primed site [14]. To identify the GSK3 sites on TRARG1, we first performed a phosphoproteomic study with GSK3 inhibitor LY2090314. As expected, phosphorylation of the known GSK3 sites on glycogen synthase (S649, S645 and S641) were significantly decreased with GSK3 inhibition (Fig. 4B). On TRARG1, S72 was the only site significantly dephosphorylated following GSK3 inhibition among the sites detected (Fig. 4B). Although phosphorylation of S80 and S76 were not detected in this study using a GSK inhibitor (Fig. 4B), they were both decreased by insulin and rescued by an AKT inhibitor (Fig. 2C), which matches the attributes of GSK3 target sites.

**Figure 4.**
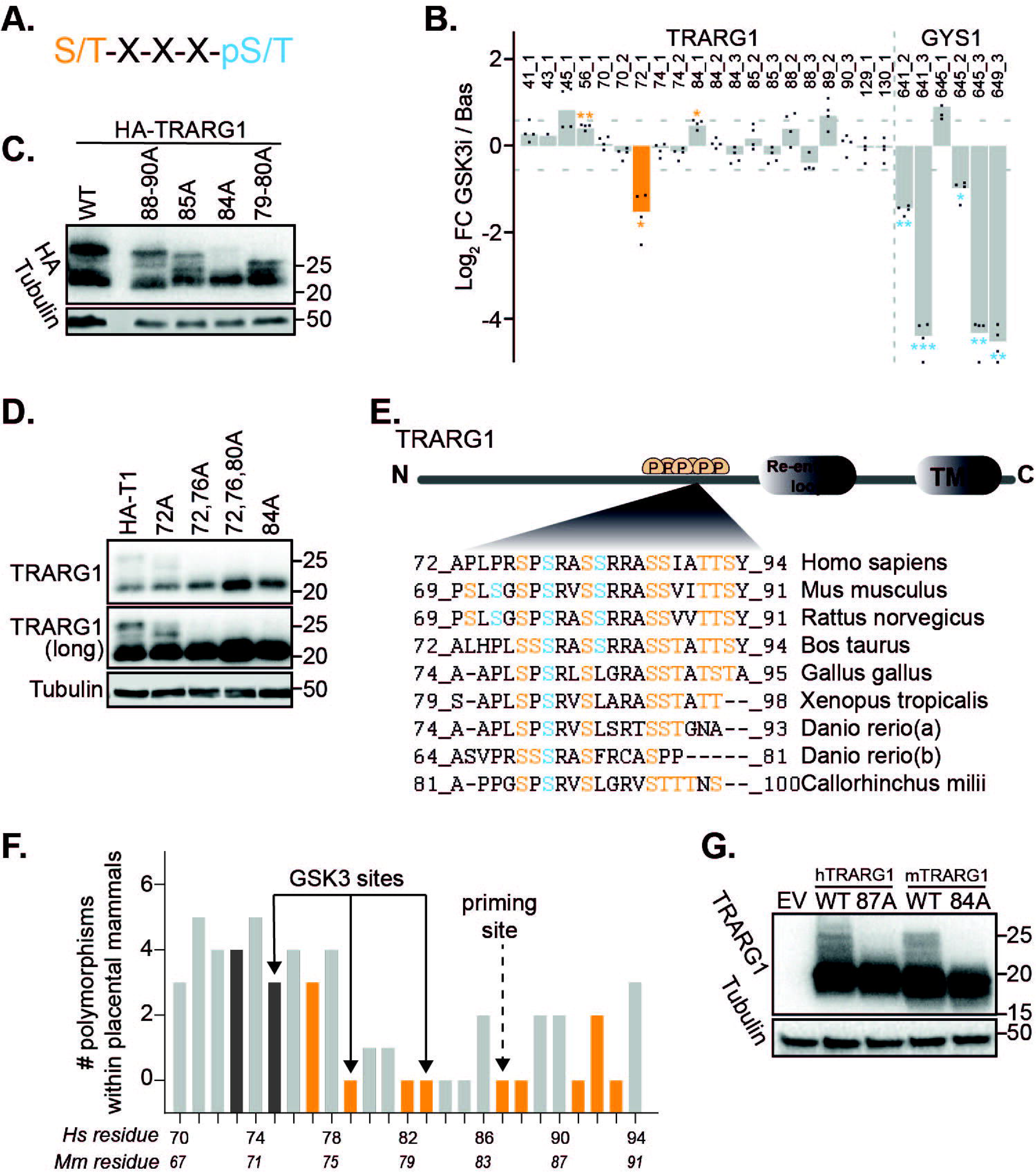
Murine TRARG1 is primed at S84 for subsequent phosphorylation by GSK3 within a highly conserved region. (A) GSK3 substrate consensus motif. Pre-phosphorylated (primed) site is labeled in *blue* and GSK3 target site is labeled in *orange*. (B) 3T3-L1 adipocytes were serum-starved followed by treatment with GSK3 inhibitor LY2090314 (100 nM, 20 min) or DMSO as a control. Cell lysates were subjected to phosphoproteomic analysis. Bar plot of log_2_-transformed mean FC of all detected TRARG1 sites and S641, S645 and S649 on glycogen synthase from this analysis is shown. Numbers following underscore indicate the number of phosphorylation sites detected on that peptide (significance is indicated by *adj. *p* <0.05; **adj. *p* <0.01; ***adj. *p* <0.001, t test). (C) Murine HA-TRARG1 phosphomutants generated by mutating T88, T89 and S90 to Ala (88-90A), S85 to Ala (85A), S84 to Ala (84A) or S79 and S80 to Ala (79-80A), or wild type HA-TRARG1 were transfected into HEK-293E cells. Cells were lysed 24 h post-transfection and samples were analyzed by immunoblotting. (D) Mouse HA-TRARG1 phosphomutants generated by mutating S72 to Ala (72A), S72 and S76 to Ala (72,76A), S72, S76 and S80 to Ala (72,76,80A) or S84 to Ala (84A), or wild type HA-TRARG1 were transfected in HEK-293E cells. Cells were lysed 24 h post-transfection and samples were analyzed by immunoblotting. A longer exposure time for the TRARG1 blot is presented to better visualize higher molecular weight bands (TRARG1 (long)). (E) The phosphosite-rich region between residue 69 and 91 on murine TRARG1 is highly conserved across vertebrate species. Insulin/GSK3 regulated sites are labeled in *blue*. Other conserved Ser/Thr residues within this region are labeled in *orange*. (F) Polymorphism of TRARG1 residues in 64 placental mammals. Ser/Thr residues conserved across murine and human TRARG1 are colored in *orange*; Bars colored in *dark grey* indicate Ser/Thr residues present in murine but not human TRARG1 sequence. Equivalent residue numbers for *mouse (Mus musculus, Mm)* and human (*Homo sapiens, Hs*) TRARG1 are shown below the bars. A full list of species included in the analysis is provided in Supplemental table S2. (G) Wild type murine TRARG1, murine TRARG1 with S84 mutated to Ala (84A), wild type human TRARG1, human TRARG1 with S87 (equivalent to S84 in murine TRARG1) mutated to Ala (87A) were expressed in HEK-293E cells. Cells were lysed 24 h post-transfection and samples were analyzed by immunoblotting. For panels C, D and G, the migration positions of molecular mass markers (kilodaltons) are shown to the right.

Therefore, these datasets together revealed a putative string of sequential GSK3 target sites on TRARG1 at T88, S84, S80, S76 and S72. Of note, S72 was not regulated by insulin in the phosphoproteomic analysis described in Fig. 2C [24] although it was clearly down-regulated in response to GSK3 inhibition (Fig. 4A).

Since phosphorylation of neither S84 nor T88 was significantly regulated by the GSK3 inhibitor, we hypothesized that S84 or S88 is the priming site (pre-phosphorylated by a different kinase). To test this, we expressed murine TRARG1 phospho-mutants with S/T at positions 79/80, 84, 85, or 88/89/90 mutated to Ala in HEK-293E cells. Only TRARG1 with S84 mutated to Ala lost the apparent higher molecular weight bands (Fig. 4C), suggesting that S84 is the priming site on murine TRARG1 allowing for its subsequent phosphorylation by GSK3 at multiple sites (S80, S76 and S72), which altered TRARG1 migration in SDS-PAGE. In support of this, murine TRARG1 with S72/76/80 mutated to Ala completely lost the apparent higher molecular weight bands, similar to the S84A mutant (Fig. 4D).

Since functionally important phosphosites are more likely conserved across species [25], we aligned TRARG1 sequences from multiple species in which TRARG1 homologues were identified. The phosphosite-rich region between residues 70 and 90 in the murine sequence is highly conserved across the vertebrate species examined (Fig. 4E; *pink* and *red* residues), indicating a likely important function for this region. Of the insulin/GSK3 regulated sites identified from phosphoproteomic analyses (Fig. 4E, *red* residues), S76 (human S79) was conserved across all species analysed, although S80 appeared to be more specific to placental mammals. Analysis of TRARG1 polymorphisms across placental mammals (Fig. 4F, Supplemental table S2) revealed that murine S76, S79, S80, S84, S85, T88 and S90 (human S79, S82, S83, S87, S88, T91 and S93) were conserved within placental mammals, while murine S72 was only present in rodents (Supplemental table S2). Indeed, the Ala mutant of S87 on human TRARG1, which is equivalent to the S84 priming site on murine TRARG1, abolished its apparent higher molecular weight bands (Fig. 4G), suggesting that this conserved site acts as the equivalent priming site for human TRARG1 phosphorylation by GSK3.

Taken together, our data show that murine TRARG1 is primed at S84 for subsequent phosphorylation by GSK3 at S80, S76 and S72. Further, these sites are highly conserved across species including human with the exception of S72, which is only present in some rodents.

### TRARG1 phosphorylation does not regulate its subcellular distribution

Since we initially observed that the apparent higher molecular weight bands of TRARG1 were highly enriched within the PM subcellular fraction (Fig. 1A), and TRARG1 undergoes insulin-stimulated translocation to the PM [5], we determined whether TRARG1 phosphorylation status altered its localization. To this end, we performed immunofluorescence microscopy with HA-TRARG1 or 7A/7E phosphomutants expressed in 3T3-L1 adipocytes. These mutants targeted the majority of the most highly conserved residues within the 70-90 phosphosite-rich region of murine TRARG1 to mitigate redundancy between phosphosites. Although the S72 and S80 sites were not mutated in this construct, the 7A or 7E mutant mimicked the lower (dephosphorylated form) or higher (phosphorylated form) band of TRARG1, respectively (Fig. 1E), likely because of the requirement for phosphorylation of S84 for GSK3-mediated phosphorylation of S72, and S80 (Fig. 4C–D). HA-TRARG1 and GLUT4 co-localized at the PM and in the peri-nuclear region (Fig. 5A), consistent with previous reports [5]. The subcellular distribution of HA-TRARG1-7A and HA-TRARG1-7E were indistinguishable from HA-TRARG1 being localized to the PM and peri-nuclear regions (Fig. 5A). These data suggest that the phosphorylation status of TRARG1 does not alter its localization.

**Figure 5.**
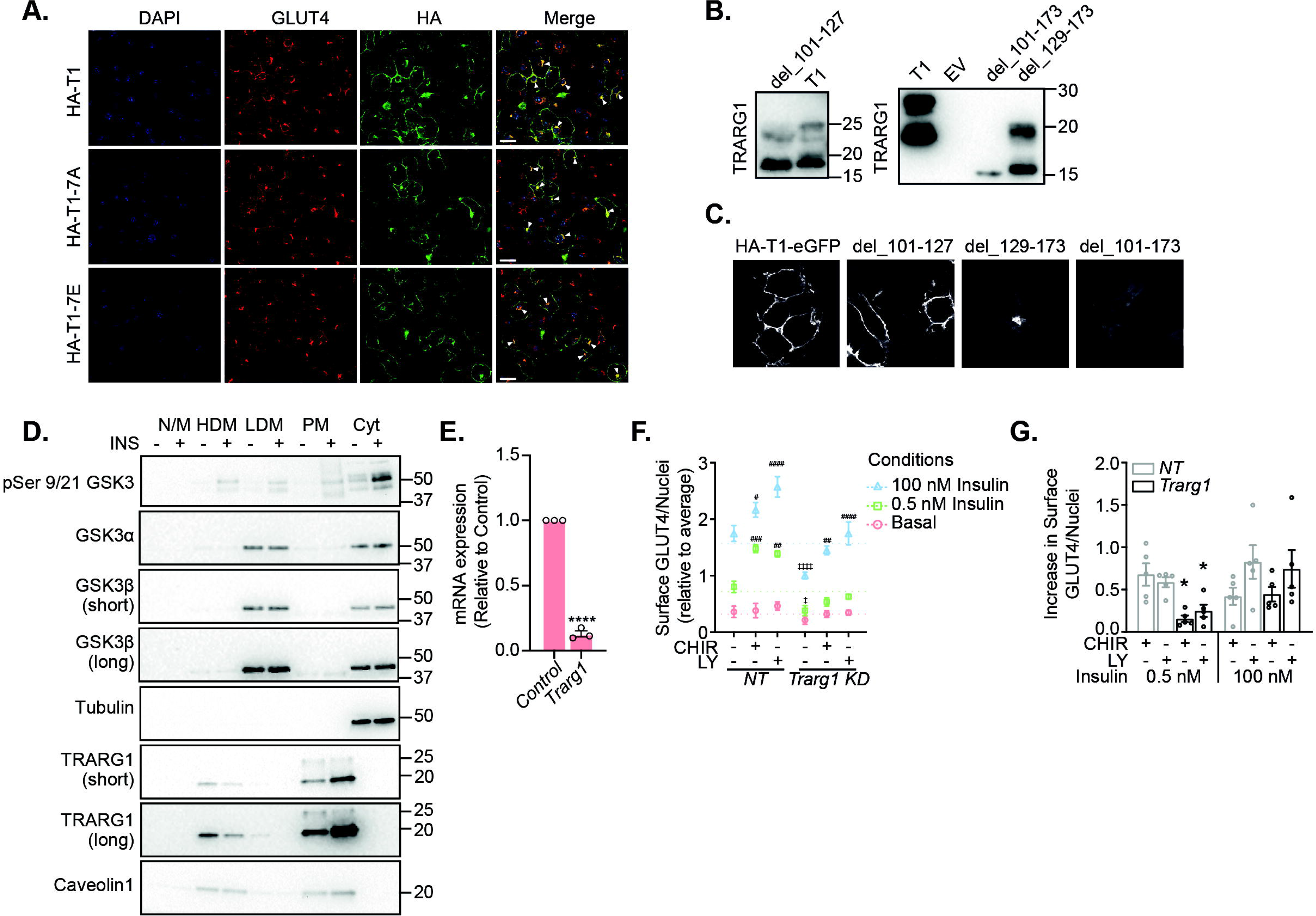
GSK3 signalling to TRARG1 does not alter TRARG1 localization but may regulate GLUT4 translocation. (A) 3T3-L1 adipocytes stably expressing HA-TRARG1, HA-TRARG1-7A or HA-TRARG1-7E were serum-starved for 2 h. Cells were fixed and stained for nuclei (DAPI, *blue*), GLUT4 (*red*) and HA (*green*). Immunofluorescence imaging was performed by confocal microscopy. Instances of colocalization between GLUT4 and HA-TRARG1, HA-TRARG1-7A or HA-TRARG1-7E are indicated by closed arrowheads. Scale bar, 20 μm. (B) Immunoblotting analysis of wild type TRARG1 (T1) or TRARG1 truncation mutants expressed in HEK-293E cells. del_101-127 and del_129-173 mutants were phosphorylated as indicated by the apparent higher molecular weight bands; del_101-173 mutant was not phosphorylated as indicated by the lack of apparent higher molecular weight band. (C) N-terminally HA-tagged and C-terminally eGFP fused TRARG1 construct (HA-TRARG1-eGFP) and its truncation mutants were expressed in HEK-293E cells. Subcellular localization of these constructs was determined by confocal microscopy. Full-length TRARG1 and del_101-127 mutant were localized to the PM; del_129-173 mutant was localized to intracellular membranes; del_101-173 mutant was cytosolic. (D) Serum-starved or insulin-stimulated 3T3-L1 adipocytes were subjected to subcellular fractionation. Subcellular localization of GSK3 was determined by immunoblotting analysis. Tubulin and caveolin1 were immunoblotted to control for loading of cytosolic and PM proteins, respectively. (E) Knockdown efficiency of *Trarg1* as assessed by qPCR (\*\*\*\**p* <0.0001, comparison with non-targeting control siRNA). (F) 3T3-L1 adipocytes were serum-starved in the absence (DMSO) or presence of GSK3 inhibitors (10 μM CHIR99021; CHIR or 500 nM LY2090314; LY) before treatment with or without 0.5 nM or 100 nM insulin for 20 min. Surface GLUT4 was quantified by immuno-labelling and expressed relative to cell number as measured by nuclei number (n=5, mean±SEM, ^#^*p* <0.05, ^##^*p* <0.01, ^###^*p* <0.001, ^####^*p* <0.0001, compared to the DMSO condition with the same insulin treatment and gene knockdown; ^‡^*p* <0.05, ^‡‡‡‡^*p* <0.0001, compared to the non-targeting (NT) knockdown with the same insulin and drug treatment). (G) Differences in PM GLUT4 between CHIR- or LY-treated and DMSO control condition with the same insulin treatment and gene knockdown as shown in (F) were calculated (mean±SEM, \**p* <0.05, compared to NT knockdown with the same insulin and drug treatment). For panel B, the migration positions of molecular mass markers (kilodaltons) are shown to the right.

As part of our previous characterization of TRARG1 topology [26], we revealed that the N-terminal cytosolic domain of TRARG1 (1-100; del_101-173) does not exhibit reduced migration by SDS-PAGE, suggesting that this truncation mutant is not phosphorylated (Fig. 5B–C). In contrast, membrane-associated truncations of TRARG1 are phosphorylated (Fig. 5B–C), suggesting that membrane localization is required for TRARG1 phosphorylation, which appears to be specific to PM-localized TRARG1 in adipocytes (Fig. 1A and 5D) [5]. In support of this, we detected GSK3 at the PM in adipocytes (Fig. 5D), and this pool of GSK3 was targeted by insulin signalling since we also detected phopsho-GSK3 in the PM fraction following insulin stimulation. These data place the kinase (GSK3) and its substrate (TRARG1) in the same subcellular location, and suggest that GSK3 at the PM is deactivated in response to insulin. Together, these data suggest that TRARG1 phosphorylation does not regulate localization, but rather TRARG1 phosphorylation is dependent on its membrane-localization.

### GSK3 signalling to TRARG1 may regulate GLUT4 translocation

Given the role of TRARG1 in insulin-stimulated GLUT4 trafficking, and previous reports that inhibition of GSK3 potentiates insulin-stimulated GLUT4 trafficking [21, 22], we tested whether TRARG1 is needed for the enhancement of insulin-stimulated GLUT4 trafficking to the PM that results from GSK3 inhibition. First, we conformed that *Trarg1* knockdown (87%; Fig. 5E) significantly decreases PM GLUT4 abundance in response to both submaximal and maximal insulin-stimulation compared to cells treated with non-targeting (NT) siRNA control (Fig. 5F), as previously reported [5, 6]. In addition, inhibition of GSK3 with either CHIR99021 or LY2090314 increased cell surface GLUT4 under insulin-stimulated conditions, compared to DMSO control cells (Fig. 5F), as suggested by previous studies using alternate GSK3 inhibitors [21, 22]. To determine whether *Trarg1* depletion blunted the effect of GSK3 inhibitors on GLUT4 trafficking, we calculated the difference in PM GLUT4 between CHIR99021- or LY2090314-treated cells and cells treated with DMSO under the same knockdown and insulin treatment conditions (Fig. 5G). The increase in PM GLUT4 in submaximal (0.5 nM) insulin-stimulated adipocytes treated with GSK3 inhibitors was attenuated by *Trarg1* knockdown (Fig. 5G). However, GSK3 inhibition increased cell surface GLUT4 to a similar extent in both control and Trarg1 KD adipocytes in response to maximal insulin stimulation (Fig. 5G). These data suggest that GSK3 activity regulates GLUT4 trafficking, and that TRARG1 is in involved in a pathway by which GSK3 inhibition increases GLUT4 trafficking to the PM at submaximal insulin doses.

## Discussion

TRARG1 colocalizes with GLUT4 and positively regulates GLUT4 trafficking and insulin sensitivity in adipocytes. However, its mechanism of action remains unclear. Here, we have integrated TRARG1 into the insulin signaling pathway and reported that TRARG1 is a novel substrate of GSK3 and its phosphorylation is regulated through the PI3K/AKT/GSK3 axis. Our data indicate that murine TRARG1 is primed by a yet-to-be-identified kinase at S84 promoting GSK3 phosphorylation of TRARG1 at S80, S76 and S72. This phosphorylation does not influence TRARG1 subcellular localization. However, *Trarg1* knockdown blunted the potentiation of submaximal insulin-stimulated GLUT4 trafficking induced by GSK3 inhibition. These data place TRARG1 within the insulin signalling pathway and suggest that GSK3 phosphorylation of TRARG1 may negatively regulate GLUT4 traffic.

The regulation of glucose transport by insulin occurs through the integration of signaling and trafficking processes that cooperate in bringing GLUT4 to the cell surface [27]. However, the precise points of intersection between upstream signaling via AKT and the distal GLUT4 trafficking machinery are not fully resolved. Although there are conflicting reports, previous studies suggested that the AKT substrate GSK3 may play a role in glucose uptake and GLUT4 traffic [15, 20-22]. However, the exact GSK3 substrates involved in this effect on GLUT4 are not yet clear. Here we corroborate a previous report that acute pharmacological inhibition of GSK3, albeit with a different GSK3 inhibitor, potentiated insulin-stimulated GLUT4 translocation (Fig. 5F) [22]. This effect was impaired by *Trarg1* knockdown at a submaximal insulin dose (Fig. 5F and G), suggesting that reduced TRARG1 phosphorylation may provide a link between GSK3 activity and GLUT4 trafficking. However, *Trarg1* knockdown did not prevent the increase in PM GLUT4 by GSK3 inhibition in cells treated with a maximal dose of insulin (Fig. 5F and G), highlighting the need for further work to understand the exact role of the GSK3-mediated TRARG1 phosphorylation in GLUT4 trafficking.

We initially observed that endogenous phosphorylated TRARG1 was enriched at the PM but not in LDM or HDM fractions [5] (Fig. 1A), suggesting that GSK3 may phosphorylate TRARG1 at this site. These data also imply that TRARG1 (and it’s phosphorylation) may play a functional role at the PM. For example, TRARG1 may affect GLUT4 internalisation or regulate interactions between GLUT4-containing vesicles and docking machinery at the PM, as reported for the TRARG1 paralogue PRRT2 [28]. Our finding that mutation of TRARG1 phosphosites did not alter TRARG1 localization (Fig. 5A), suggests that localization may impart phosphorylation status, and not *vice versa*. Indeed, GSK3 has been reported to localize to the PM at the early stage of WNT signaling activation [29] and our subcellular fractionation analysis showed that GSK3 was mainly localized to the LDM and cytosol, with a small proportion of GSK3 localized to the PM (Fig. 5D). Further supporting a role for TRARG1 localization in promoting phosphorylation, the TRARG1 mutant (del_101-173), which abolishes its PM/membrane localization, was not phosphorylated in HEK-293E cells as indicated by the lack of higher molecular weight bands (Fig. 5B–C). Of note, a TRARG1 mutant (del_129-173) localized to intracellular membranes was still phosphorylated in HEK-293E cells, suggesting that there may be differential control of TRARG1 phosphorylation between HEK-293E cells and adipocytes. Thus, our data suggest that TRARG1 phosphorylation on residues between 70 and 90 does not affect TRARG1 localization, but rather TRARG1 phosphorylation is localization-dependent.

TRARG1 amino acid sequence alignment across multiple species revealed a highly conserved phosphosite-rich region between residues 70 and 90 in the murine TRARG1 N-terminal cytosolic region, with conservation shared by *Callorhincus milii* (Australian ghost shark), the earliest extant species in which TRARG1 is found. Using changes in phosphosite abundance in response to insulin and GSK3 inhibition and phosphosite mutagenesis, we identified a GSK3-priming site (murine S84) and GSK3 phosphorylation sites (murine S72, S76 and S80). The S76 and S80 GSK3 target sites were highly conserved, at least within placental mammals (Fig. 4F), suggesting an important role of regulated phosphorylation for TRARG1 function. One key difference within mammals is the presence of a third GSK3 site in rodents (murine S72), that is almost exclusively proline in human and other mammals (Supplemental table S2). Nevertheless, the loss of phosphorylation upon mutation of the S87 priming site in human TRARG1 (Fig. 4G) and loss of endogenous human TRARG1 phosphorylation in SGBS adipocytes treated with a GSK3 inhibitor (Fig. 3G) suggest that our findings using murine TRARG1 are translatable to human TRARG1.

Hotspots of protein phosphorylation such as those we have identified in TRARG1 have been implicated in modulating protein-protein interactions [25], and may serve as an integration point for multiple signaling pathways [30]. For example, changes in TRARG1 phosphorylation status may release or recruit regulators of GLUT4 traffic. Indeed, in general there were multiple apparent higher molecular weight immuno-reactive TRARG1 bands observed in both HEK cells and adipocytes (Fig. 1H, 3A–C, E, H–K), suggesting distinct phospho-species. These TRARG1 phospho-species depend on GSK3 since inhibition or knockdown of GSK3 almost completely ablated all apparent higher molecular weight forms of TRARG1 (Fig. 3B and K). This suggests the multiple apparent higher molecular weight bands result from phosphorylation at some of the GSK3 sites, but not others (e.g. S80 and S76, but not S72), or that GSK3-mediated phosphorylation of TRARG1 leads to phosphorylation at alternate sites not targeted directly by GSK3 (Fig. 1C and 4A). Understanding the role that distinct TRARG1 phospho-species play in TRARG1 function will be the focus of future studies. In addition, our findings that TRARG1 is a new GSK3 substrate raises several questions including: 1) the identity of the priming kinase for subsequent GSK3 activity; 2) whether other kinases can also target this region of TRARG1 in addition to GSK3; and 3) which phosphatase dephosphorylates TRARG1.

In summary, we have integrated the trafficking regulator of GLUT4-1, TRARG1, into the insulin signaling network via GSK3 and provided evidence that TRARG1 may be involved in the mechanisms by which GSK3 inhibition promotes insulin-stimulated GLUT4 trafficking. This provides strong basis for future studies into the exact mechanisms by which GSK3-to-TRARG1 signaling regulates GLUT4 trafficking and insulin sensitivity in adipocytes.

### Experimental procedures

#### Molecular Cloning

pcDNA3.1-HA-TRARG1, pcDNA3.1-TRARG1 and pcDNA3.1-HA-TRARG1-eGFP constructs were generated as previously described [26]. HA-TRARG1 sequences containing 7 (S74, S76, S79, S84, S85, T88, S90) or 12 (S70, S72, S74, S76, S79, S80, S84, S85, T88, T89, S90, Y91) Ser/Thr/Tyr to Ala point mutations and 7 or 11 (S70, S72, S74, S76, S79, S80, S84, S85, T88, T89, S90) Ser/Thr to Glu point mutations were generated from pcDNA3.1-HA-TRARG1, using Gibson assembly cloning [31, 32]. These constructs were designated as pcDNA3.1-HA-TRARG1-7A, pcDNA3.1-HA-TRARG1-12A, pcDNA3.1-HA-TRARG1-7E and pcDNA3.1-HA-TRARG1-11E, respectively. These sequences were then shuttled into pBABE-puro vector using Gibson assembly cloning. Using overlap-extension site-directed mutagenesis [33], we mutated all the Cys in HA-TRARG1 sequence into Ser, which resulted in the construct pcDNA3.1-HA-TRARG1-C-S. Using Gibson assembly cloning, we mutated the first four Lys at the N-terminus of HA-TRARG1 into Arg, followed by the mutagenesis of the last three Lys into Arg using overlap-extension site-directed mutagenesis, the resulting construct was designated as pcDNA3.1-HA-TRARG1-K-R (see Fig. 1D for details of Ser/Thr/Tyr, Lys and Cys mutants used in this study). TRARG1 truncation mutants pcDNA3.1-HA-TRARG1-del_129-173-eGFP and pcDNA3.1-HA-TRARG1-del_101-173-eGFP were generated as previously described [26]. Residues 101-127 were deleted from pcDNA3.1-HA-TRARG1-eGFP using overlap-extension site-directed mutagenesis to yield the plasmid pcDNA3.1-HA-TRARG1-del_101-127-eGFP. pcDNA3.1-TRARG1-del_101-127, pcDNA3.1-TRARG1-del_129-173, and pcDNA3.1-TRARG1-del_101-173 were generated from pcDNA3.1-TRARG1 by deleting residues 101-127, 129-173, or 101-173, respectively, using overlap-extension site-directed mutagenesis. Primer sequences are available upon request. Constructs were confirmed by Sanger sequencing.

#### Cell culture and transfection

3T3-L1 fibroblasts and HEK-293E/HeLa cells were maintained in Dulbecco’s modified Eagle’s medium (DMEM) supplemented with 10% fetal bovine serum (FBS) and GlutaMAX (Life Technologies, Inc.) (DMEM/FBS medium) at 37 °C with 10% CO_2_. L6 myoblasts were grown in α-MEM supplemented with 10% FBS and GlutaMAX at 37 °C and 10% CO_2_. For 3T3-L1 stable cell lines, fibroblasts were transduced with pBABE-puro retrovirus (empty vector control), or retrovirus with the constructs of interest in pBABE-puro vector. Puromycin (2 μg/mL) was used to select for transduced cells. 3T3-L1 fibroblasts were cultured and differentiated into adipocytes as described previously [5], and used for experiments between day 9 and 12 after the initiation of differentiation. HEK-293E/HeLa cells were transfected with indicated constructs 2 days before experiments using lipofectamine2000 (Thermo Scientific), according to the manufacturer’s instructions. Myoblasts at 80% confluence were differentiated into myotubes by replacing 10% FBS with 2% FBS. Myotubes were used in experiments at between 6-8 days post-differentiation. SGBS cells were cultured and differentiated as previously reported [34, 35], and used for experiments 12-14 days after initiation of differentiation. SGBS adipocytes were incubated in growth-factor free media for 4 h prior to addition of insulin or GSK3 inhibitors.

#### Cell fractionation

3T3-L1 adipocytes were washed with ice-cold PBS and harvested in ice-cold HES-I buffer (20 mM HEPES, pH 7.4, 1 mM EDTA, 250 mM sucrose containing protease inhibitors mixture (Roche Applied Science)). All subsequent steps were carried out at 4 °C. Cells were fractionated as previously described [26]. Briefly, cells were homogenized by passing through a 22-gauge needle 5 times and a 27-gauge needle 10 times prior to centrifugation at 500 x g for 10 min to pellet cell debris. The supernatant was centrifuged at 13,500 x g for 12 min. The pellet contained the PM and mitochondria/nuclei (M/N). The supernatant consisted of cytosol, low density microsome (LDM), and high density microsome (HDM). This supernatant was centrifuged at 21,170 x g for 17 min to pellet the HDM fraction. The supernatant was again centrifuged at 235,200 x g for 75 min to obtain the LDM fraction (pellet). The pellet containing PM and M/N was referred to as PM fraction in this study as TRARG1 is not enriched in M/N fractions (data not shown).

### Lambda protein phosphatase assay in 3T3-L1 adipocytes/HEK-293E and fat tissues

Cells were washed with PBS twice and lysed in 1% (v/v) Triton X-100 in PBS and then solubilized by passing through a 22-gauge needle 3 times and a 27-gauge needle 3 times prior to centrifugation at 12,000 x g for 15 min. In the case of fat tissues, subcutaneous/epidydimal white adipose tissue or brown adipose tissue was excised from mice and lysed in 1% (v/v) Triton X-100 in PBS. Tissues were lysed and homogenized by sonication (Gallay Scientific) prior to centrifugation at 13,000 x g for 15 min. The supernatant was transferred to a clean tube and protein concentration determined by bicinchoninic acid (BCA) assay (Thermo Scientific). 100 μg of tissue or cell lysates were treated with or without 2 μL Lambda Protein Phosphatase (LPP) (New England Biolabs) at 30 °C for 15 min (30 min for HEK-293E lysate) in a reaction volume of 50 μL. Starting material without 15 min (30 min for HEK-293E lysate) incubation at 30 °C was included as a control. Reaction was terminated by the addition of SDS (1% (w/v) final concentration), and the reaction mixture was incubated at 95 °C for a further 10 min. Loading sample buffer (LSB) with TCEP was added to the samples, which were then incubated at 65 °C for 5 min, separated by SDS-PAGE, and analyzed by immunoblotting.

#### Phosphatase inhibitors assay and kinase inhibitors assay in 3T3-L1 adipocytes

3T3-L1 fibroblasts were seeded in 12-well plates and differentiated into adipocytes. Cells were used for experiments on day 10 post-differentiation. For phosphatase inhibitors assay, adipocytes were treated with 100 nM calyculin A, 1 μM okadaic acid or DMSO in DMEM/FBS medium for 60 min. For kinase inhibitors assay, cells were serum-starved in basal DMEM media (DMEM, GlutaMAX, 0.2% (w/v) bovine serum albumin (BSA, Bovostar, Bovogen)) for 2 h prior to treatment with kinase inhibitors (100 nM wortmannin, 10 μM MK-2206, 100 nM rapamycin or 1 μM GDC-0994) for 20 min followed by 100 nM insulin treatment for 20 min. Cells were transferred onto ice and washed with ice-cold PBS twice, followed by lysis with 1% (w/v) SDS in PBS-containing protease inhibitors mixture (Roche Applied Science) and phosphatases inhibitors mixture (1 mM sodium pyrophosphate, 2 mM sodium orthovanadate, 10 mM sodium fluoride). Cell lysates were sonicated at 90% intensity for 12 s and centrifuged at 13,000 x g. Protein concentration of the supernatant was determined by BCA assay. LSB was added to the samples, which were then incubated at 65 °C for 5 min, separated by SDS-PAGE, and analyzed by immunoblotting.

#### GSK3 inhibitors treatment in adipose tissue explants

Epididymal or subcutaneous fat depots were excised from mice, transferred immediately to warm DMEM/2% BSA/20 mM HEPES, pH 7.4, and minced into fine pieces. Explants were washed 3 times and incubated in DMEM/2% BSA/20 mM HEPES, pH 7.4 for 2 h. Explants were then aliquoted and treated with 10 nM insulin, 500 nM LY 2090314 or DMSO for 30 min at 37 °C. Treatment was terminated with three rapid washes in ice-cold PBS, after which the cells were solubilized in RIPA buffer (50 mM Tris-HCl, pH 7.5, 150 mM NaCl, 1% (v/v) Triton X-100, 0.5% (w/v) sodium deoxycholate, 0.1% (w/v) SDS, 1 mM EDTA, protease inhibitors mixture). Samples were then subjected to sonication prior to spin at 20,000 x g for 15 min at 4 °C. The supernatant was transferred to a clean tube and protein concentrations were determined using BCA assay. LSB with TCEP was added to the samples, which were then incubated at 65 °C for 5 min, separated by SDS-PAGE, and analyzed by immunoblotting.

#### EsiRNA-mediated GSK3 Knockdown

EsiRNA (Merck, EMU170441, EMU059761) was added to 100 μL Opti-MEM medium to a final concentration of 900 nM with 7.5 μL transfection reagent TransIT-X2 (Mirus Bio), mixed and incubated at room temperature (RT) for 30 min. Adipocytes at 6 days post-differentiation were washed once with PBS, trypsinized with 5 x trypsin-EDTA (Life Technologies, Inc.) at 37 °C, resuspended in DMEM/FBS medium and then centrifuged at 200 x g for 5 min. The supernatant was removed, and pelleted cells were resuspended in DMEM/FBS medium of the same volume as that of the media where the cells were previously cultured (e.g. 1 mL of media for the cells from one well of a 12-well plate). 900 μL of cell suspension was then reseeded onto one well of a Matrigel-coated 12-well plate. 100 μL of esiRNA or luciferase control mixture was added into per well of a 12-well plate (esiRNA final concentration 90 nM). Cells were used in experiments 72 h following esiRNA knockdown. Method relevant to Fig. 3K–M.

#### Immunoprecipitation of TRARG1 and in vitro kinase assay

3T3-L1 adipocytes at day 10 post-differentiation were serum-starved in basal DMEM media containing 100 nM LY2090314 for 2 h. Cells were then transferred to ice, washed thrice with ice-cold PBS and harvested in lysis buffer (1% NP 40, 5% glycerol, 50 mM Tris-HCl, pH 7.4, 150 mM NaCl) containing protease inhibitors mixture and phosphatase inhibitors mixture. Cells were lysed by passing through a 22-gauge needle six times, followed by six times through a 27-gauge needle. Lysates were solubilized on ice for 20 min and then centrifuged at 20,000 x g for 20 min at 4 °C to remove insoluble material. The protein concentration of the supernatant was quantified by BCA assay following the manufacturer’s protocol. For each sample, 2.5 mg protein was combined with 5 μL of anti-TRARG1 mouse monoclonal antibody (200 μg/mL) (sc-377025) or 2.5 μL of mouse anti-IgG (400 μg/mL) in a final volume of 500 μL and incubated with rotation at 4 °C for 1 h. Magnetic Dynabeads (Life Technologies, 10004D) were washed once with ice-cold PBS, once with lysis buffer and then resuspended in lysis buffer. 50 μL of magnetic beads were then added into each immunoprecipitation (IP) tube followed by incubation with rotation at 4 °C for 1 h. Beads were then separated from flow-through by magnetic capture and resuspended and washed twice with lysis buffer, followed by two washes with ice-cold PBS. The residual liquid was removed from the final wash. The TRARG1-IP tubes and IgG control tube were incubated with either kinase assay buffer (20 mM Tris-HCl, pH 7.4, 10 mM MgCl_2_, 1mM DTT, 1 mM sodium pyrophosphate and 10 mM sodium fluoride, 10 mM glycerol-2-phosphate, 2 mM ATP) or kinase assay buffer containing either 1 μg GSK3α or 1 μg GSK3β in a final volume of 30 μL at 37 °C for 1 h. Following the incubation, reactions were terminated, and proteins were eluted by the addition of 30 μL 2X laemmli sample buffer and incubation at 65 °C for 5 min. Samples were then centrifuged at 20,000 x g for 2 min at RT and stored at −20 °C.

#### Polymorphism analysis

TRARG1 sequences were extracted from Ensembl (release 99) and aligned by Multiple sequence Alignment using Fast Fourier Transform (MAFFT) version 7 [36]. Analysis of the number of polymorphisms at each residue was conducted across 64 unique placental mammalian species (represented by 69 genomes including different subspecies/sexes of the same species). The results of this analysis between residues 67-91 of murine TRARG1 are presented in Fig. 4F and the full analysis in Supplemental table S2.

#### Immunofluorescence confocal microscopy

3T3-L1 adipocytes stably expressing HA-TRARG1, HA-TRARG1-7A or HA-TRARG1-7E were re-plated onto Matrigel-coated glass coverslips (Matrigel from Becton Dickinson) on day 6 after initiation of differentiation and processed for immunofluorescence microscopy on day 10. Cells were washed with PBS and fixed with 4% (v/v) paraformaldehyde (PFA) for 20 min at RT. Cells were blocked and permeabilized with 2% (w/v) BSA and 0.1% (w/v) saponin in PBS followed by incubation with a mixture of rabbit anti-HA antibody (CST, C29F4) and anti-GLUT4 1F8 antibody (generated in-house) at RT for 45 min. Cells were washed five times with 2% (w/v) BSA and 0.1% (w/v) saponin in PBS. Anti-mouse Alexa-555-conjugated secondary antibody (Life Technologies) and anti-rabbit Alexa-488-conjugated secondary antibody (Life Technologies) were used to detect primary antibodies. DAPI was used to visualize nuclei. Samples were mounted in Immuno-Fluore Mounting Medium (MP Biomedicals). For HEK-293E cells expressing HA-TRARG1 truncation mutants, cells were re-plated onto Matrigel-coated glass coverslips in 12-well plate 24 h after transfection, allowed to adhere overnight. Samples were prepared as described above, but without antibody incubations. Optical sections were obtained using Nikon C2 Confocal microscope using a Plan Apo VC 60X WI DIC N2 (NA=1.2 WD=270 μm) objective. Images were acquired using NIS Elements software. All images were processed using Fiji [37].

#### siRNA-mediated gene knockdown for GLUT4 translocation assays

47 μL of OptiMEM, 2 μL of TransIT-X2 and 1 μL of siRNA (25 μM, ON TARGETplus siRNA pool (Dharmacon, L-057822-01 for *Trarg1*, D-001810-10 for non-targeting pool), final concentration 500 nM) were mixed and incubated for 30 min at RT. On day 6 post-differentiation, 3T3-L1 adipocytes grown in a 6-well plate (stably expressing HA-GLUT4-mRuby3 where appropriate) were washed once with PBS, trypsinized with 5X trypsin-EDTA at 37 °C, resuspended in DMEM/FBS medium and centrifuged at 200 x g for 5 min. The supernatant was removed, and pelleted cells were resuspended in 13.5 mL DMEM/FBS. 450 μL of cell suspension was added into the mixture of OptiMEM/TransIT-X2/siRNA and mixed well (final concentration of siRNA was 50 nM). All 500 μL of the cell suspension plus OptiMEM/TransIT-X2/siRNA was added to one well of a Matrigel-coated 24-well plate, or 84 μL of the cell suspension plus OptiMEM/TransIT-X2/siRNA was aliquoted into each well of the Matrigel-coated 96-well plate. Cells were refed with DMEM/FBS medium 24 h post transfection and used 72 h post transfection. Method relevant to Fig. 5F–G.

#### Endogenous GLUT4 translocation assay

Cells were washed once with PBS and once with basal DMEM media prior to incubation in basal DMEM media for 2 h in a CO_2_ incubator. Cells were stimulated with 0.5 nM or 100 nM insulin for 20 min. For endogenous GLUT4 translocation assay (Fig. 5F–G), cells were serum-starved in the presence or absence of DMSO (control) or GSK3 inhibitor prior to 0.5 nM or 100 nM insulin stimulation for 20 min. After the stimulation, cells were washed by gently dunking the 96-well plates 12 times in a beaker containing ice cold PBS (all the subsequent PBS washes were performed using this method). Plates were placed on ice and residual PBS was removed with multichannel pipette. Cells were fixed with 4% PFA for 5 min on ice, and 20 min at RT and PFA was replaced with 50 mM glycine in PBS followed by incubation for 15 min. After dunking the plates 6 times in RT PBS, residual PBS was removed and cells were blocked with 5% Normal Swine Serum (NSS, Dako, X0901) in PBS for 20 min. After removing all the blocking media, cells were incubated with human anti-GLUT4 antibody (LM048; [38]) (Integral Molecular, PA, USA) in 2% NSS in PBS for 1 h to detect GLUT4 present at the PM. After 12 washes in RT PBS, residual PBS was removed and cells were incubated with anti-human Alexa-488 (Life Technologies) and Hoechst 33342 (Life Technologies) in 2% NSS in PBS for 1 h. Cells were washed 12 times in RT PBS and stored in PBS containing 2.5% DABCO, 10% glycerol, pH 8.5 and imaged on the Perkin Elmer Opera Phenix High Content Screening System. Imaging data were analyzed using Harmony Software supplied with the imaging system. GLUT4 signal was normalized to the number of nuclei per imaging region (as measured by Hoechst 33342 signal).

#### Sample preparation and real-time quantitative-PCR assays

For Fig. 5E, total RNA was extracted from cells using QIAshredder and RNeasy kits (Qiagen). To remove any DNA, the extracts were incubated with DNAse buffer (Promega) and residual DNAse was subsequently inactivated with DNAse stop solution (Promega). cDNA synthesis was performed using LunaScript RT SuperMix Kit (NEB). Polymerase chain reactions were carried out using TaqMan 2X Universal PCR Master Mix or SYBR Green PCR Master Mix (Thermo) on a QuantStudio™ 7 Flex Real-Time PCR System (Thermo). Acidic ribosomal phosphoprotein P0 (36B4), β-actin (b-act) and 18S ribosomal RNA (18s) were used as internal controls. The following primer sets were used: 36B4_F; AGATGCAGCAGATCCGCAT and 36B4_R; GTTCTTGCCCATCAGCACC, b-act_F; GCTCTGGCTCCTAGCACCAT and b-act_R; GCCACCGATCCACACAGAGT, and 18s_F; CGGCTACCACATCCAAGGAA and 18s_R; GCTGGAATTACCGCGGCT, with the corresponding 18s taqman probe GAGGGCAAGTCTGGTGCCAG. The TaqMan gene expression assay (premixed primer set and probes) was used for murine *Trarg1* (Mm03992124_m1).

#### Western blotting

Typically, 10 μg of protein was resolved by SDS-PAGE, transferred to PVDF membranes and immunoblotted as previously described [5]. Antibodies detecting TRARG1 (sc-292062 or sc-377025) and 14-3-3 (sc-1657) were from Santa Cruz Biotechnology. Antibodies against pHSL S660 (#4126), pp90RSK (#9344), p4EBP1 S65 (#9456), pTBC1D4 T642 (#4288S), pAKT S473 (#4051), pAKT T308 (#9275L), AKT (#4685), HA (#C29F4), GSK3α (#9338S), GSK3β (#9832S), pGSK3 Ser 9/21 (#8566S), pGS S641 (#3891) and Caveolin1 (#3267) were purchased from Cell Signaling Technology. Anti-α-tubulin (#T9026) was from Sigma-Aldrich. Antibody against TBC1D4 was produced as previously described [4].

#### Sample preparation for MS-based analsyis of TRARG1

HEK-293E cells transiently expressing HA-TRARG1 or TRARG1 were washed three times with ice-cold PBS, lysed in RIPA buffer and homogenized prior to centrifugation at 20,000 x g, 4 °C for 20 min to remove cellular debris. 2 mg of each cell lysate was incubated with 40 μL Anti-HA Microbeads (μMACS HA Isolation Kit, Miltenyi Biotec, 130-091-122) in a volume of 500 μL for 45 min at 4 °C with rotation. Microbeads were separated from flow-through by running through μColumns (MACS, Miltenyi Biotec, 130-042-701) and three washes with RIPA buffer, followed by two washes with ice-cold PBS. Proteins were eluted in 2X laemmli sample buffer and subjected to SDS-PAGE. Gels were stained with SyproRuby protein gel stain (Life Technologies) according to the manufacturer’s instructions and imaged on a Typhoon FLA 9500 biomolecular imager (GE Healthcare). TRARG1 bands were excised for in-gel digestion.

Gel fractions were washed twice in 50% acetonitrile (ACN, Thermo Scientific) in 100 mM ammonium bicarbonate (NH_4_HCO_3_, Sigma) at RT for 5min at 2,000 rpm in a ThermoMixer C (Eppendorf). Liquid was removed, 10 mM TCEP and 40 mM 2-Chloroacetamide (Sigma) in 100 mM NH_4_HCO_3_ was added to the gel fractions and incubated at RT for 30 min at 2,000 rpm. Liquid was removed and gel pieces were dehydrated in 100% ACN at RT for 5 min at 2,000 rpm followed by rehydration in 100 mM NH_4_HCO_3_ containing 14 ng/μL trypsin (Promega) on ice for 1 h. Excess liquid was removed and 100 μL of 100 mM NH_4_HCO_3_ was added to gel slices and samples were incubated at 37 °C overnight. Peptides were solubilized by spiking in Trifluroacetic acid (TFA, Thermo Scientific) to 1% and incubating at 37°C for 30 min. To extract peptides, gel pieces were dehydrated using 100% ACN and incubated at 37°C for 15 min. Peptides were transferred to a new tube and dried in a vacuum concentrator (Eppendorf) and then resuspended and acidified in 1% TFA.

Resuspended peptides were centrifuged at 21,000 x g for 10 min at 4°C. The peptide mixture was desalted using stage-tips containing styrene divinylbenzene-reverse phase sulfonated (SDB-RPS) (Empore, 3M). StageTips were pre-equilibrated with addition of 100% ACN, 30% methanol (Fisher Scientific) and 1% TFA then 0.2% TFA and 5% ACN and centrifugation at 1,500 x g for 3 min. After loading the samples, StageTips were washed once with 99% isopropanol/1% TFA and once with 0.2% TFA/5% ACN, then peptides were eluted from StageTips with elution buffer (60% ACN, 5% NH4OH (20% v/v of a 25% NH_4_OH solution)). Samples were dried in a vacuum concentrator for ~45□min, at 45□°C and resuspended in 5_μl MS loading buffer (0.3% TFA, 2% ACN) for MS analysis.

For phosphoproteomics analysis in 3T3-L1 adipocytes treated with GSK3 inhibitor (Fig. 4B), samples were lysed and prepared using the recently described EasyPhos method [39, 40], before being resuspended in 5□μL MS loading buffer (2% ACN, 0.3% TFA).

#### Mass spectrometry

For analysis of immunoprecipitated TRARG1, an Easy nLC-1000 UHPLC was connected to a Q Exactive mass spectrometer for mass spectrometry analysis. A 75□μm□×□40□cm column packed in-house (ReproSil Pur C18-AQ, 1.9□μm particle size) was used to separate the peptides prior to elution with a gradient of 5–30% ACN containing 0.1% formic acid (FA) at 200□nL/min over 90□min with the column temperature set at 55□°C. Peptides were analyzed with one MS scan from 300 to 1750□m/z with a resolution of 70,000, 3e^6^ AGC, and a max injection time (IT) of 100□ms, followed by data-dependent MS/MS scans with HCD at a resolution of 17,500, AGC 5e^5^, IT 60□ms, NCE 25%, and isolation width of 2.0□m/z. For phosphoproteomics analysis in 3T3-L1 adipocytes treated with GSK3 inhibitor, label-free quantification was applied and samples measured on a Q Exactive HF-X mass spectrometer (Thermo Fisher Scientific) [41] as previously described [42].

#### Processing of spectral data

Raw mass spectrometry data was processed using the Andromeda algorithm integrated into MaxQuant (v1.6.6.0 or v1.6.1.0) [43], searching against the mouse UniProt database (June 2019 release) concatenated with known contaminants. Default settings were used for peptide modifications, with the addition of Phospho(STY) for the phosphoproteomics study, or the addition of Phospho(STY), GlyGly(K), Oxidation(M), Acetyl(K), Deamidation(NQ), Methyl(KR) in the variable modifications for immunoprecipitated TRARG1. Match between runs was turned on with a match time window of 0.7 min and an alignment time window of 20 min for identifications transfer between adjacent fractions, only for samples analyzed using the same nanospray conditions. For immunoprecipitated TRARG1, only murine TRARG1 sequence was used for searching. Protein, peptide and site FDR was filtered to 1% respectively.

#### Quantification and statistical analysis

##### GSK3 inhibitor phosphoproteomic study

This study was performed with four biological replicates. Phosphopeptides groups were filtered out for reverse sequences, potential contaminants and those only quantified 2 times or fewer out of the 8 samples. LFQ intensities were log_2_-transformed and median normalized. Each group was imputed as previously described in [44]. For remaining missing values, a second step of imputation was performed if at least three replicates had quantified values in one condition and the values were completely missing or had only one quantified replicate in the other condition, using a method previously described in [45]. For the phosphopeptides with no missing values after imputation, two sample t tests were performed. Treatment was compared to control and p values were corrected for multiple hypothesis testing using the Benjamini and Hochberg method [46].

##### Western blotting intensities

In phosphatase inhibitor assays, differences between control and treated intensities were tested with unpaired two-sided student’s t-tests with Welch’s correction. In kinase inhibitor assays, differences were tested by one-way ANOVA with Dunnett’s multiple comparisons test. In tissue explants experiment, differences were tested with paired two-sided t-tests. In GSK3 knockdown experiments, differences were tested by RM one-way ANOVA with Holm-Sidak’s multiple comparisons test. Analyses were performed using GraphPad Prism version 8.0 for macOS (GraphPad Software, CA USA). Error bars are presented as standard error of the mean (SEM). Significance is represented, with a p-value < 0.05 by *, < 0.01 by **, < 0.001 by *** and < 0.0001 by ****.

##### Endogenous GLUT4 translocation assays

For endogenous GLUT4 translocation assay, experimental values were normalized to the average value across all conditions in each experiment. Differences of surface GLUT4 relative to nuclear DNA among different conditions were tested using two-way ANOVA with correction for multiple comparisons. Differences of increase of surface GLUT4 relative to nuclear DNA among different conditions were tested using paired two-sided t-tests. Analyses were performed using GraphPad Prism version 8.0 for macOS (GraphPad Software, CA USA). Error bars are SEM. Significance is represented as stated in figure legends.

### Data Access Statement

Raw and MaxQuant processed data of MS-based proteomics (except for the phosphoproteomic analysis of GSK3 inhibitor treated cells, which will be uploaded to accompany a future publication) have been deposited in the PRIDE ProteomeXchange Consortium (http://proteomecentral.proteomexchange.org/cgi/GetDataset) [47] and can be accessed with the identifier PXD022765.

## Acknowledgements

We thank Dr Joseph Rucker and Integral Molecular (PA, USA) for kindly providing the LM048 anti-GLUT4 antibody and Professor Martin Wabitsch (Division of Pediatric Endocrinology and Diabetes, University Medical Center Ulm, Germany) for generously providing SGBS cells. These studies were supported by Sydney Mass Spectrometry, the Wellcome-MRC, Institute of Metabolic Science, Metabolic Research Laboratories, Imaging Core (Wellcome Trust Major Award [208363/Z/17/Z]).

## Funding and additional information

X.D. was supported by China Scholarship Council (CSC)-The University of Sydney (USYD) joint postgraduate scholarship (201606270221). W.P.B. and F.M.B. were supported by Wellcome Trust Investigator Award (107858/Z/15/Z) and UKRI-MRC grant MR/S008144/1 to F.M.B. B.L.P was supported by a National Health and Medical Research Council (NHMRC) Early Career Fellowship APP1072129. O.J.C was supported by a Wellcome Trust PhD studentship in Basic Science. J.G.B. was supported by a Diabetes Australia Research Program grant. J.R.K. was supported by a Diabetes Australia Research Program grant and an Australian Diabetes Society Skip Martin Early-Career Fellowship. D.E.J was supported by a NHMRC Senior Principal Research Fellowship APP1019680. D.J.F was supported by a Diabetes Australia Research Program grant and UKRI-MRC Career Development Award MR/S007091/1. The contents of the published material are solely the responsibility of the authors and do not reflect the views of the NHMRC.

## Rights Retention Statement

This work was funded by UKRI grant MR/S007091/1. For the purpose of open access, the author has applied a Creative Commons Attribution (CC BY) licence to any Author Accepted Manuscript version arising.

---

**Supplementary table S1.** Identified post-translational modifications on murine Trarg1.

**Supplementary table S2.** Analysis of *TRARG1* conservation and polymorphisms in placental mammals.

